# Resolving Taxonomic Conflicts in Trichiurus: Chromosome Genomics Clarifies Species and Informs Conservation of Populations in the Northwest Pacific

**DOI:** 10.1101/2025.03.05.641545

**Authors:** Vanthu Giap, Tianqin Wu, Liang Lu, Jiantao Hu, Mingwei Zhou, Chenhong Li

**Author notes:** Contributed equally to this work (Vanthu Giap, Tianqin Wu). Author for correspondence: Chenhong Li.

## Abstract

The Trichiuridae family, comprising ecologically and commercially important hairtails, has faced persistent taxonomic uncertainties due to historical misclassifications and conflicting morphological criteria. While molecular approaches have advanced understanding, mitochondrial-based phylogenies and misapplied nomenclature have obscured species boundaries and population dynamics. Here, we integrate a chromosome-level genome assembly of *Trichiurus japonicus* with whole-genome resequencing of congeneric species to resolve three centuries of taxonomic ambiguity. Combining phylogenomic, demographic, and selective sweep analyses, we resolve taxonomic uncertainties by redefining species boundaries, reconstructing population dynamics, and identifying genomic adaptations underpinning environmental resilience. We confirm *T. japonicus* and *T. nanhaiensis* as distinct species. Crucially, we identify a cryptic Indian Ocean-derived lineage, misclassified as *T. lepturus* in previous studies, as the novel species *Trichiurus* sp., underscoring the impact of commercial trade on taxonomic confusion. Population genomics reveals panmixia in *T. japonicus* across the Northwest Pacific, supporting unified stock management. Selective sweeps in *T. japonicus* pinpoint cold-resilient genes (metabolic, anti-apoptotic, visual), consistent with its demographic stability during Pleistocene glaciations. This study provides a genomic roadmap for disentangling taxonomic conflicts in morphologically cryptic taxa and advances strategies for sustainable fisheries management under climate change.

## Introduction

The Trichiuridae family, comprising ecologically and commercially critical species commonly termed hairtails, ribonfish or cutlassfishes, sustains global fisheries with annual yields exceeding 1.2 million metric tons (FAO 2020). In the Northwest Pacific, ten species are documented, though only three—*Trichiurus japonicus*, *T. lepturus*, and *T. nanhaiensis*, hold commercial significance (Wang et al., 2017). Effective fisheries management relies on precise species delineation and population structure data, yet morphological convergence among taxa obscures these parameters. While molecular markers have advanced genetic characterization of fishery species (Abdul-Muneer, 2014), persistent taxonomic conflicts in *Trichiurus* hinder sustainable management strategies.

Taxonomic ambiguities in *Trichiurus* trace to Linnaeus’ 1758 misclassification of three morphologically similar taxa under *T. lepturus*. Subsequent revisions identified *T. japonicus* from Japanese specimens (Temminck & Schlegel, 1843) and *T. haumela* from East China Sea specimens (Chu, 1931), yet mitochondrial analyses later confirmed *T. haumela* as a junior synonym of *T. japonicus* (Yi et al., 2022). Despite this, misapplication of invalid nomenclature (e.g., *T. haumela* in Liu et al., 2021; Jin et al., 2013) and erroneous classification of *T. japonicus* as *T. lepturus* (Panhwar et al., 2018; Kim et al., 2011) persist, underscore the need for systematic resolution to inform conservation strategies. Furthermore, all existing phylogenies of these species are mitochondrial-based, which may not fully reflect nuclear genomic divergence. Thus, chromosome-level genome assemblies and genome-wide SNP analyses are urgently needed to reconcile these taxonomic inconsistencies.

Wang et al. (1991) first described *T. nanhaiensis* and *T. brevis* from South China Sea specimens, though current NCBI database entries and primary research frequently misclassify *T. nanhaiensis* as the subspecies *T. lepturus nanhaiensis*. Concurrently, Li et al. (1992) proposed *T. margarites* (now a junior synonym of *T. nanhaiensis*) for morphologically comparable specimens. We have collected both *T. lepturus* and *T. nanhaiensis* from the South China Sea, enabling genomic evaluation of their species status. Yamada (1995) subsequently identified an undescribed taxon (*T.* sp.2), with debates regarding its specific versus subspecific relationship to *T. margarites*. Mitochondrial analyses clustered *T. nanhaiensis* and *T.* sp.2 within a monophyletic clade, suggesting potential conspecificity (Guo et al., 2012). We also discovered a cryptic *Trichiurus* species (*T.* sp) from the fish markets in China and Korea, showing highest cytochrome c oxidase subunit I gene (COXI) sequence similarity to Indian *T. lepturus* but clustering with *T. nanhaiensis* in preliminary phylogenetic reconstructions, which may represent the unresolved *T.* sp.2 lineage.

Population delineation is critical for fisheries management, but remains equally contentious in *Trichiurus* in the Northwest Pacific. Mitochondrial haplotype analyses propose eight distinct *T. japonicus* stocks in the East China Sea (Wen, 2024), while others recognize only East and South China Sea stocks (Tzeng, C. H., 2016). Historical overexploitation of Yellow Sea spawning aggregations during 1950-1960 reportedly caused local extirpation of *T. japonicus*. Subsequent analyses by Wang et al. (1995) identified migrant individuals from Zhoushan populations in Haizhou Bay specimens post 1970, though no eggs were detected during 1980 spawning surveys. By the 1990s, emerging genetic differentiation between Haizhou Bay and Zhoushan stocks suggested partial population recovery. Our comprehensive sampling across Chinese (Qingdao, Xiamen, Zhoushan, Shanghai, Dandong, Hainan) and South Korean (Busan) coastal waters aims to clarify contemporary population structure and inform adaptive management strategies.

This study integrates chromosome-level genome assembly of *T. japonicus* and comparative whole-genome resequencing of it and congener species to resolve the centuries old taxonomic inconsistency. By integrating phylogenomic, demographic, and selective sweep analyses, we address species boundaries, glacial-era population dynamics, and adaptive genomic signatures, providing a framework for sustainable fisheries management in the Northwest Pacific.

## Result

### Chromosome-Level Genome Assembly and Functional Annotation

We generated a high-quality chromosome-level assembly of the *Trichiurus japonicus* genome through integration of 112 Gb Oxford Nanopore long reads, 66.37 Gb Illumina short reads, and 104.11 Gb Hi-C scaffolding data. The final assembly comprised 24 chromosomes (1,038 scaffolds; 964.99 Mbp total length) with 41.21% GC content and scaffold/contig N50 values of 39.19 Mbp and 6.07 Mbp, respectively (Table 1). Genome annotation, supported by 58.91 Gb of transcriptomic data, identified 23,993 protein-coding genes.

**Table 1.** Summary of genome assembly of *Trichiurus japonicus*.

| Assembly | Data |
| --- | --- |
| Total length (Mbp) | 964.99 |
| GC (%) | 41.21 |
| Scaffold N50 length (Mbp) | 39.19 |
| Contig N50 length (Mbp) | 6.07 |
| Scaffold N50 number | 1038 |
| Contig N50 number | 1825 |
| Gene number | 23993 |
| Heterozygosity (%) | 1.2 |
| Total BUSCO (%) | 95.8 |
| Repeat content (%) | 40.48 |
| Hi-C anchoring rate (%) | 98.87 |
| Number of chromosomes | 24 |
| Percent gaps (%) | 0.082 |

Repetitive elements constituted 40.48% of the genome, dominated by retroelements (10.50%; primarily LINEs [3.89%] and LTRs [6.10%]), DNA transposons (10.95%), and unclassified repeats (14.08%). Genome quality assessments revealed 96.9% BUSCO completeness (95.8% single-copy genes, 1.1% duplicated). GenomeScope 2.0 analysis indicated a homozygous genome (98.8%) with minor heterozygosity (1.2%).

Comparative synteny analysis demonstrated strong chromosomal conservation between *T. japonicus* and related Perciformes (*Scomber japonicus*, *Thunnus albacares*), with collinear blocks spanning entire chromosomes (Fig. 2A). Genomic feature mapping revealed spatial correlations between syntenic regions, GC content gradients, gene density hotspots, and transposable element distributions (Fig. 2B).

**Fig. 1.**
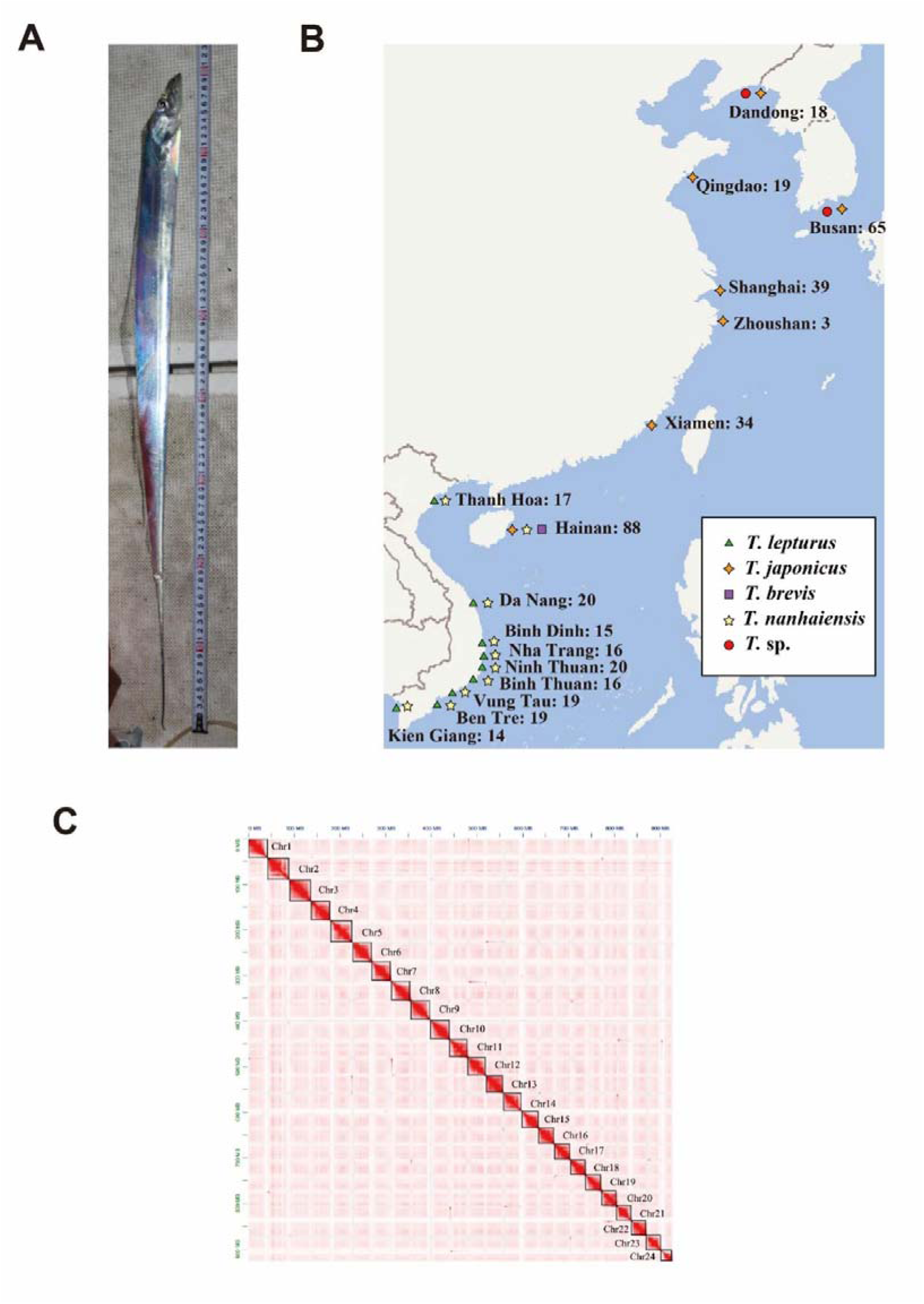
**A**. The specimen of *Trichiurus japonicus* used for genome sequencing, collected by hand fishing rod in Zhoushan, China **B**. Sampling locations for genome resequencing. **C**. Chromosome contact map of *T. japonicus*.

**Fig. 2.**
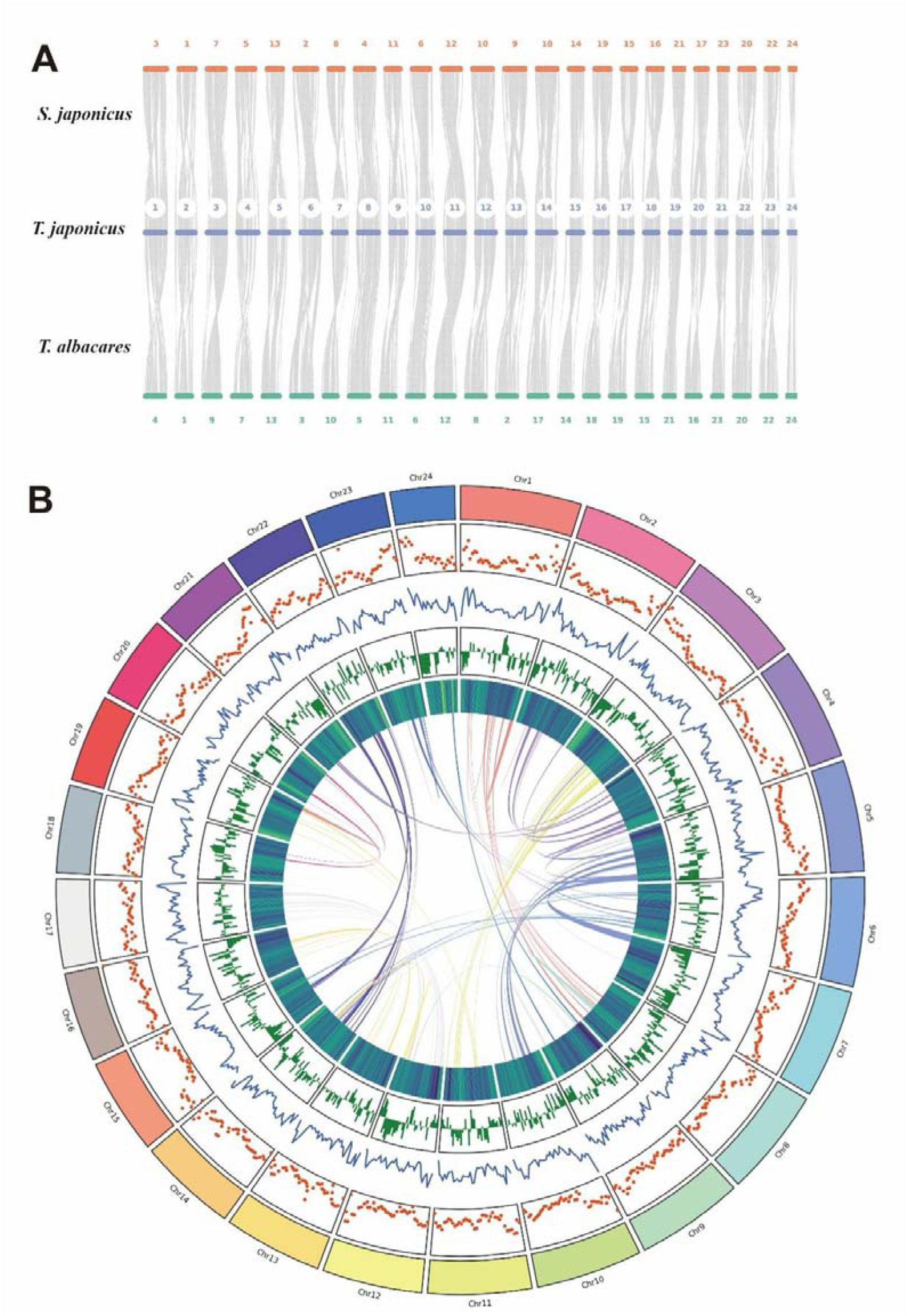
**A**. Synteny plot showing the genomic collinearity between the *Trichiurus japonicus*, *Scomber japonicus* and *Thunnus albacares*. **B**. Genomic features of the *T japonicus*. From the inner to the outer layer: genome-wide synteny, GC content heatmap, gene density distribution, LTR transposon distribution, and TE transposon distribution

### Population genomics analysis

Principal Component Analysis (PCA) revealed clear genetic divergence among *Trichiurus* taxa across the Northwest Pacific (Fig. 3A). *Trichiurus japonicus* formed a tight cluster (PC1: 49.02%, PC2: 44.65% variance explained), indicative of limited genomic diversity and a stable evolutionary trajectory. *Trichiurus lepturus* showed phylogenetic isolation along PC1, confirming deep divergence from congeners. In contrast, *T.* sp. and *T. nanhaiensis* exhibited overlapping confidence ellipses (Procrustes similarity = 0.9), suggesting genomic similarity, while PC2 distinguished *T. japonicus* from the *T.* sp.-*T. nanhaiensis* complex.

**Fig. 3.**
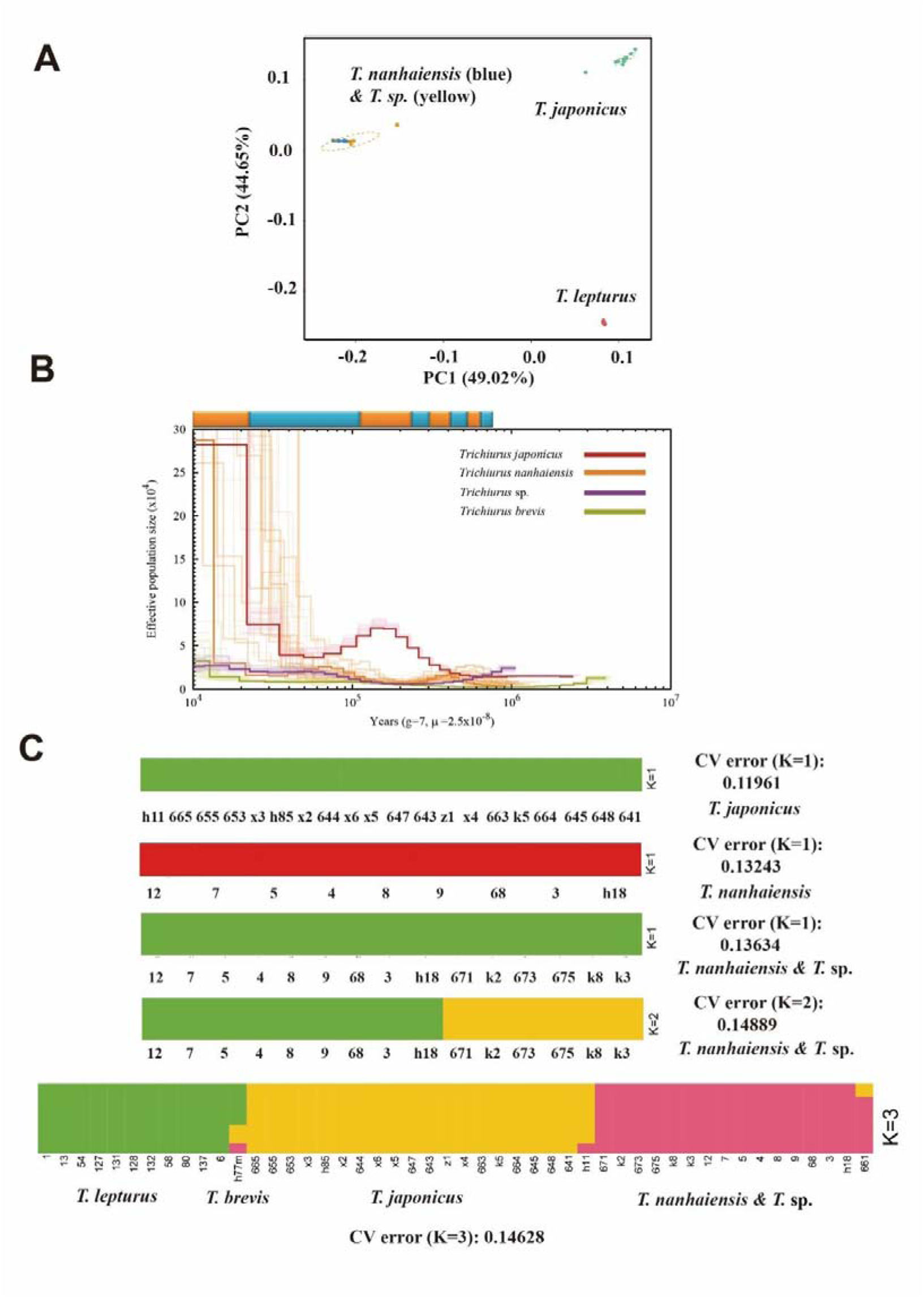
**A**. Principal Component Analysis (PCA) of four *Trichiurus* taxa. **B**. Demographic history of *Trichiurus* lineages reconstructed via Pairwise Sequentially Markovian Coalescent (PSMC) analysis (100 bootstrap replicates). The timeline bar denotes glacial (blue) and interglacial (orange) periods. **C**. Admixture proportions for five *Trichiurus* taxa, inferred using genome-wide SNPs.

Admixture analysis revealed that *T. japonicus* samples from South Korea, Hainan, Zhoushan, and Xiamen formed a single genetic population, indicating genetic homogeneity. *Trichiurus nanhaiensis* exhibited a similar lack of population subdivision across Hainan and Vietnamese waters. In contrast, *T.* sp. clustered with *T. nanhaiensis* in PCA and admixture analyses (Fig. 3C), suggesting their taxonomic closeness. Admixture analysis of *Trichiurus nanhaiensis* and *Trichiurus* sp. revealed minimal cross-validation error at K=1, yet distinct genetic clusters emerged at K=2. Subsequent phylogenetic analyses demonstrated reciprocal monophyly between these two lineages, rejecting hypotheses of their conspecificity. These results instead satisfy the primary species criterion of non-overlapping genealogical exclusivity, warranting their classification as distinct biological species.

Admixture analyses identified a single genetic cluster of *T. japonicus* across South Korea, Hainan, Zhoushan, and Xiamen, supporting panmixia. Similarly, *T. nanhaiensis* showed no subdivision between Hainan and Vietnam. However, *T. sp.* clustered with *T. nanhaiensis* in PCA but formed distinct genetic groups in admixture analyses (Fig. 3C). Admixture analysis of *Trichiurus nanhaiensis* and *Trichiurus* sp. revealed minimal cross-validation error at K=1, yet distinct genetic clusters emerged at K=2. Subsequent phylogenetic analyses demonstrated reciprocal monophyly between these lineages, rejecting conspecific hypotheses. This genealogical exclusivity meets the primary species criterion, supporting their classification as distinct species.

### Phylogenomic analysis

Phylogenomic reconstruction resolved distinct evolutionary lineages within *Trichiurus* (Fig. 4A). *T. lepturus* and *T.* sp. formed reciprocally monophyletic clades, with a basal lineage comprising South China Sea *T. lepturus* specimens. Notably, *T. brevis* exhibited discordant placement: mitochondrial phylogenies positioned it outside the *T. lepturus* complex, while nuclear markers placed it intermediate between *T. lepturus* and *T. japonicus*. Admixture analysis revealed hybrid ancestry in *T. brevis* (59.1% *T. lepturus*, 26.1% *T. japonicus*, 14.7% *T. nanhaiensis*), suggesting historical introgression (Fig. 3C). Mitochondrial genome assemblies independently confirmed the distinct status of *T.* sp. (Fig. 4B).

**Fig. 4.**
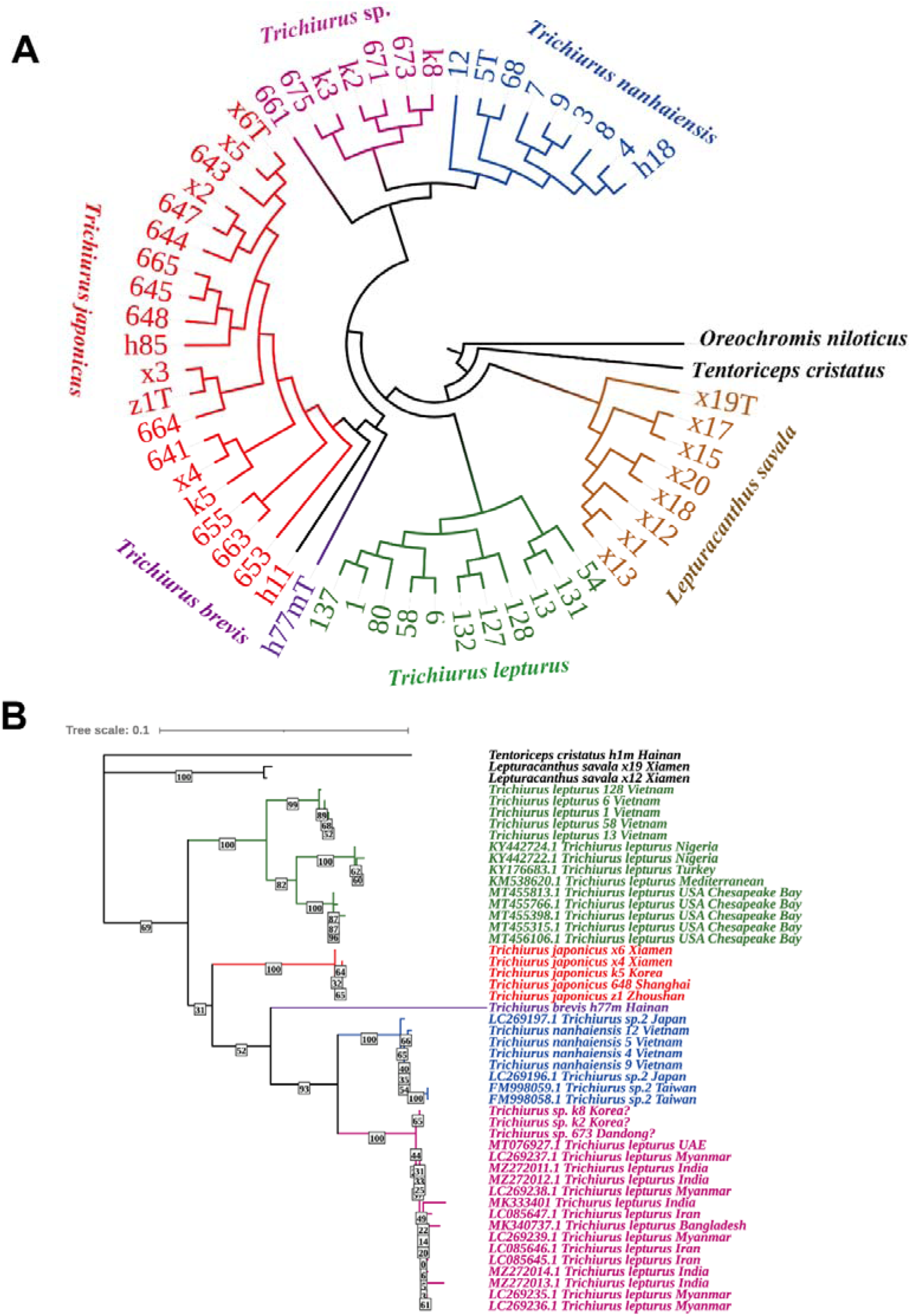
**A**.Phylogenetic relationship among resequencing samples of *Trichiurus* reconstructed using Raxml. **B** Phylogenetic relationship of Trichiurus based on the COX1 sequences extracted from resequencing data and the downloaded data from the different regions of the world.

Our findings will demonstrate that the previously misidentified as endemic hairtail specimens were in fact imported from the Indian Ocean, while simultaneously revealing a persistent taxonomic misclassification within the *Trichiurus* genus across the whole Indian Ocean marine region. Comprehensive genetic analyses confirm reveal that previous studies on *Trichiurus* from the Persian Gulf to the Bay of Bengal have systematically misidentified a distinct species as *T. lepturus*, highlighting critical flaws in current ichthyological taxonomy of Indian Ocean hairtail populations.

### Demographic History and Divergence Timing

Using 100 independent Fastsimcoal2 bootstrap simulations, we reconstructed demographic parameters—including population sizes, divergence times (in generations), and gene flow patterns—for four *Trichiurus* species (Fig. 5A). *Trichiurus japonicus* exhibited the largest effective population size. The optimal model inferred a divergence sequence beginning with *T. japonicus* splitting from the *T. lepturus* ancestor 200,875 generations ago, followed by *T. nanhaiensis* diverging from *T. japonicus* 11,712 generations ago, and finally *T.* sp. separating from *T. nanhaiensis* 11,522 generations ago. Contemporary gene flow among extant species was negligible.

**Fig. 5.**
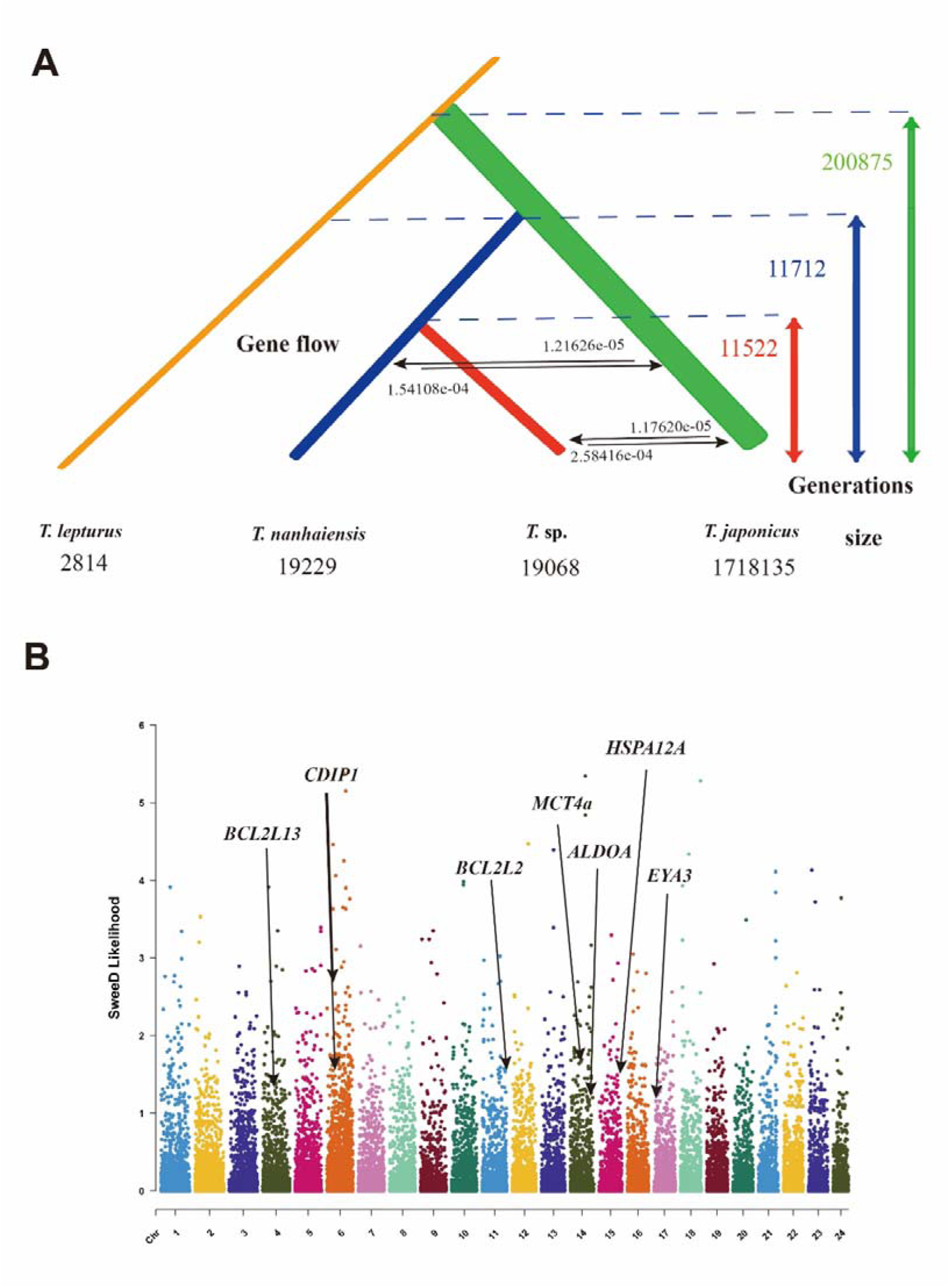
**A**. Demographic history of four *Trichiurus* species reconstructed from 100 Fastsimcoal2 bootstrap simulations, depicting effective population sizes (Ne), divergence times (generations), and gene flow patterns. **B** Genomic signatures of selection in *T. japonicus* identified via SweeD analysis, showing loci under positive selection (top 1% composite likelihood ratio [CLR] scores) and associated cold-adaptive genes.

Assuming a generation time of 1–3 years (based on rapid sexual maturity in hairtails), the *T. nanhaiensis* split dates to ∼11,000–33,000 years before present (YBP), coinciding with Holocene marine transgressions. The earlier *T. nanhaiensis* speciation aligns with the Last Dali Glaciation (20,000–70,000 YBP; Hermann & Wissmann, 2002), when sea-level regression in the East and South China Seas exposed continental shelves, creating biogeographic barriers that drove reproductive isolation from *T. lepturus*.The *T. japonicus* lineage emerged earlier (200,000–500,000 YBP), corresponding to the Lushan Glacial Period (Li, 1947). PSMC analyses corroborated this timeline, showing sustained large Ne in *T. japonicus* but negligible ancestral signals in *T. nanhaiensis* and *T.* sp., consistent with their more recent origins.

### Genomic Signatures of Cold Adaptation

SweeD analysis identified genomic regions under selection (top 1% likelihood scores) in *T. japonicus*, revealing key cold-resilience adaptations (Fig. 5 B). Selective sweeps targeted ALDOA (fructose-bisphosphate aldolase A, a glycolytic enzyme) and MCT4a (monocarboxylate transporter 4a, a lactate shuttle), likely optimizing anaerobic metabolism during glacial cooling in the East China Sea. Cold-induced p53 pathway activation was inferred from selection on CDIP1 (cell death-inducing p53-target protein 1) and anti-apoptotic genes (BCL2L13, BCL2L2), potentially reducing temperature-dependent cell death. Additionally, HSPA12A (heat shock protein 70 family) may mitigate cold-induced protein misfolding through enhanced repair mechanisms. Enhanced visual perception in low-light glacial habitats is suggested by selection on eya3 homologue (eye development), improving foraging efficiency. These coordinated genetic changes underscore *T. japonicus*’s evolutionary resilience to Pleistocene glacial cycles, balancing metabolic efficiency, cellular survival, and ecological performance.

## Discussion

### Genomic Architecture and Evolutionary Insights

The chromosome-level assembly of *T. japonicus* (964.99 Mbp) provides unprecedented resolution for comparative genomics within Trichiuridae. The genome structural conservation—evidenced by size similarity to congeners (*T. nanhaiensis*: 868 Mb; *T. brevis*: 871 Mb) and stable GC content (39.59–42.05%), suggests strong evolutionary constraints buffering genomic architecture against speciation-driven divergence. High continuity metrics (scaffold N50: 39.19 Mbp; contig N50: 6.07 Mbp) and BUSCO completeness (96.9%) underscore its utility for macroevolutionary studies. With 23,993 protein-coding genes, *T. japonicus* surpasses *Eupleurogrammus muticus* (21,949 genes) but aligns closely with *Lepturacanthus savala* (23,625 genes), reflecting conserved regulatory frameworks despite ecological divergence. Synteny with *Scomber japonicus* and *Thunnus albacares* underscores shared perciform ancestry, further validating the assembly’s phylogenetic utility.

### Resolving The Centuries’ Taxonomic Discordance

Linnaeus’ 1758 misclassification of multiple lineages under *T. lepturus* initiated persistent taxonomic confusion. A previous study by Li et al. (2006) on Linnaeus’s type specimens proved that *T. lepturus* sensu Linnaeus should be regarded as a nomen dubium, as its original description (*protologue*) confounds four diagnosable species spanning three extant genera *(Trichiurus, Lepturacanthus, and Eupleurogrammus*). Subsequent revisions, such as Temminck & Schlegel’s (1844) *T. japonicus*, clarified species boundaries but were undermined by synonymization errors. For instance, *T. haumela*, an invalid epithet misapplied to *T. japonicus* by Asian taxonomists, originated from Forsskål’s (1775) ambiguous type specimens, which lack definitive ties to Northwest Pacific populations. European taxonomists predominantly regard *T. haumela* as a junior synonym of *T. lepturus*, (Golani, D., & Fricke, R., 2018) a conclusion stemming from Forsskål’s original problematic designation. Conversely, Asian researchers erroneously synonymized *T. japonicus* with *T. haumela* (Wang et al., 1995), a misconception traceable to Chu’s (1931) misidentification of *T. japonicus* and subsequent inappropriate application of the *haumela* epithet. We therefore advocate formal rejection of *T. haumela* to rectify historical inconsistencies.

### Misclassification on Imported Specimens

Wang’s (1991, 1992) description of *T. nanhaiensis* and *T. brevis* from South China Sea specimens, alongside Li’s (1992) concurrent proposal of synonymous taxa *T. margarites* and *T. minor*, generated further taxonomic discord. Wang’s (1995) methodological critique of Li’s work exposed factual inaccuracies and flawed analytical approaches, though both agreed on the existence of only two valid species. Notably, Wang identified Chu’s 1931 classification as the root cause of modern *T. haumela* confusion. Subsequent taxonomic consensus has validated Wang’s *nanhaiensis* and *brevis* classifications.

Guo et al. (2012) acknowledged persistent international inconsistencies in *lepturus* delineation while advocating retention of *T. margarites*. Their mitochondrial analysis of Japanese *T.* sp.2 specimens, which was firstly classified by Nakabo (2002), morphologically similar yet purportedly differentiated by premaxillary tubercle size, clustered *T.* sp.2 with *T. margarites*, separating them from *T. japonicus* and *T. lepturus* clades.

Our investigation involved with the acquisition of hairtail specimens (*Trichiurus* sp.) from markets in Dandong, China and Busan, South Korea, where vendors claimed their local origin. Initially hypothesized to represent a potential Yellow Sea population, subsequent analysis prompted suspicions of an Indian Ocean origin. This conclusion is substantiated by the following evidence.

Genome-wide phylogenetic reconstruction revealed that *T.* sp forms a monophyletic clade with hairtail specimens from the South China Sea, displaying greater genetic divergence from *T. lepturus* than from *T. japonicus*. Mitochondrial COI sequence analysis, conducted using NCBI BLAST showed 99% similarity with “*T. lepturus*” specimens from Indian Ocean collections (Mukundan et al., 2020, MK333401). Additionally, comparative analysis with *T.* sp.2-Nakabo specimens (Okamoto et al. 2018, LC219626, LC219697) revealed clustering within South China Sea hairtails, suggesting taxonomic alignment with this group.

Previous studies utilized imported hairtail specimens from Qingdao’s market as outgroups, which similarly clustered with Indian specimens MK333401 (Li et al. 2022). While the precise geographic origin of the *T.* sp specimen remains uncertain, the consistency between our findings and those reported by the referenced authors warrants careful consideration. Specifically, their specimen QDM8 was conclusively identified as imported hairtail (distinct from Chinese domestic varieties) and formed a monophyletic clade with Indian specimens (MK333401,collected from Munambam, Kochi, along the southwestcoast of Indian Ocean), while maintaining a sister-clade relationship with *T. nanhaiensis* (as phylogenetically closest yet distinct sister clades). Our data exhibit striking parallels, leading us to propose that the *T.* sp specimen in question likely represents a distinct species of imported hairtail, with its geographic origin strongly indicating biogeographic affinity to the Indian Ocean.

Expanded phylogenetic analysis, incorporating COI sequences from specimens in the UAE, Iran, India, Bangladesh, and Myanmar, revealed consistent clustering of Indian Ocean specimens. Furthermore, this COI gene-based phylogenetic tree reveals that the monophyletic group from the Indian Ocean is genetically distant to the true *T. lepturus* populations originating from Turkey, the Americas, West Africa, and the South China Sea, and therefore should not be classified as the same species. Given that the genetic misclassification of *T. lepturus* in the Indian Ocean has been substantiated, the conclusion designating *T. nanhaiensis* as a subspecies of *T. lepturus* rather than a separate species appears increasingly untenable. Phylogeny also strongly supported elevating *T. nanhaiensis* to full species status (Fig. 4 B), with reciprocal monophyly in both mitochondrial (COX1) and nuclear markers—a critical distinction absent in previous studies limited to single-marker systems. *Trichiurus* sp. formed a distinct clade sister to *T. nanhaiensis*, potentially representing a cryptic species adapted to the Indian Ocean condition, though expanded sampling across the Bohai Sea is required.

Hybrid ancestry in *T. brevis* indicated historical admixture involving *T. lepturus*, *T. japonicus*, and *T. nanhaiensis*, requiring further genomic and ecological investigation.

### Demographic Resilience and Fisheries Implications

Pairwise Sequentially Markovian Coalescent (PSMC) analysis resolved distinct demographic trajectories among Trichiuridae species (Fig. 3B). During the Dali Glacial Period (57–16 kya), *T.* sp., *T. nanhaiensis*, and *T. brevis* maintained low effective population sizes (0–2). In contrast, *T. japonicus* retained high pre-glacial population estimates (>6), experiencing <30% reductions during glacial maxima. PSMC trajectories revealed differential glacial responses within the genus. *Trichiurus japonicus* maintained stable effective population size during the Dali Glacial Period (MIS 3-4, 30-60 kya), corroborating its ecological dominance as evidenced by historical catch records exceeding 700,000 tons annually in the East China Sea. Conversely, severe bottlenecks in *T.* sp. and *T. nanhaiensis* align with Last Glacial Maximum habitat fragmentation, suggesting lower thermal tolerance thresholds. This demographic resilience difference explains *T. japonicus*’ current biogeographic dominance across subtropical-temperate transition zones.

ADMIXTURE analysis (K=1) confirmed panmixia in *T. japonicus* across the Northwest Pacific, supporting its management as a single fishery stock—a critical consideration given its status as China’s highest-yielding marine fish. The anomalous clustering of “*T. lepturus* (*T.* sp.)” to *T. nanhaiensis* indicates persistent taxonomic misidentification, necessitating reclassification as *T.* sp.

### Taxonomic Revisions and Database Errors

*Trichiurus* spp. is the most consumed marine fish in China, consistently leading annual capture yields over the past decade (China Fishery Statistical Yearbooks). However, recent mitochondrial genome studies on Chinese coastal populations revealed inconsistent phylogenetic relationships due to specimen misidentification and ambiguous taxonomy. Taxonomic revisions and phylogenetic assessments of *Trichiurus* remain scarce, leading to mislabeling in public databases. For instance, Li et al. (2022) misidentified a specimen (likely *T.* sp.) as *T. lepturus* due to reliance on a misannotated *T. lepturus* mitochondrial genome (NCBI: MK333401), which our COX1 phylogeny placed within *T. nanhaiensis*. This finding aligns with the mitochondrial phylogeny of Yi et al. (2022), wherein *T. nanhaiensis* from South China Sea and *Trichiurus lepturus* specimens from Indian waters formed a monophyletic clade in gene tree reconstructions.

Further errors include Wen (2023) misclassifying NCBI sequence MW719078 as *T. japonicus*, inferring eight implausible populations in Chinese waters. Jiao (2024) misidentified *T. japonicus* as *T. haumela* (MH846121) and retained the outdated name *T. lepturus nanhaiensis* (NCBI: MW719076), which molecular data supports as *T. nanhaiensis*. These misclassifications stress the need for taxonomic standardization to prevent errors in fisheries management.

### Outlook and Limitations

The hybrid ancestry detected in *T. brevis* (59.1% *T. lepturus*, 26.1% *T. japonicus*) requires validation through expanded sampling, particularly in putative parental habitats. This conclusive evidence awaits integrative genomic and morphological analyses.

Our findings offer crucial insights for future global surveys of hairtail (*Trichiurus* spp.) fishery resources. The results underscore the methodological necessity of direct onboard sampling over reliance on local market specimens to ensure traceability of biological origins - though such convenient sampling practices remain common in marine fisheries research. The unexpected inclusion of Indian Ocean-derived *T.* sp. specimens in our analyses and other studies introduced significant taxonomic confusion, initially suggesting a putative new species in the Yellow Sea. This revelation fundamentally challenges the conventional species delineation within the *Trichiurus* genus, compelling a revision of the existing taxonomic framework based on genetic evidence. However, limitations in specimen acquisition and preservation protocols currently hinder comprehensive morphological validation. Future research directions should prioritize coordinated international sampling efforts to obtain sufficient voucher specimens, enabling formal morphological description and further taxonomic resolution.

## Conclusion

This study resolves centuries’ of taxonomic discordance in *Trichiurus* through integrative phylogenomics, validating *T. japonicus* and *T. nanhaiensis* as distinct species and identifying the cryptic lineage from Indian Ocean (*T.* sp.) as a novel species. By combining chromosome-level genome assembly and comparative resequencing, we reconcile mitochondrial phylogenetic conflicts, underscoring the necessity of genome-scale data for disentangling complex speciation histories. Population genomic analyses reveal *T. japonicus*’s demographic stability during Pleistocene glaciations, driven by selection in cold-resilience genes, while *T. nanhaiensis* and *T.* sp. experienced severe bottlenecks due to lower thermal tolerance. The panmictic structure of *T. japonicus* across the Northwest Pacific mandates unified fishery management. Crucially, our findings reveal cryptic speciation within commercial hairtail populations in specimens collected from Chinese and Korean markets, identifying as a genetically distinct lineage of Indian Ocean origin misidentified as *T. lepturus*. Our phylogenetic analyses including sequences retrieved from NCBI and documentation of database mislabels underscore the urgent need for updated bioinformatic resources. Future efforts must prioritize direct specimen sampling and international collaboration to disentangle biogeographic complexities, because the taxonomic status of *T.* sp. requires formal morphological diagnosis and broader biogeographic sampling before formal species description. This integrative approach bridges taxonomic clarity with adaptive genomics, providing a blueprint for sustaining marine fisheries under escalating climatic and anthropogenic pressures.

## Materials and Method

### Sample collection and sequencing

One specimen of *T. japonicus* (Fig. 1 A) was captured via handline fishing near Gouqi Island, Shengsi County, Zhoushan City, Zhejiang Province, China (geographic coordinates: 30°N, 122°E) on 23 October 2023. Liver tissue was immediately flash-frozen in liquid nitrogen for Nanopore sequencing, while muscle tissue was preserved for Hi-C library construction. Ten additional tissue types (heart, liver, brain, kidney, skin, eye, muscle, swim bladder, gill, and blood) were stabilized in RNA later solution for transcriptomic analyses.

Genomic DNA extraction from hepatic tissue followed the CTAB protocol using the QIAGEN Genomic DNA Extraction Kit (#13343). For Nanopore sequencing, 3-4 μg of high-molecular-weight DNA was processed into libraries and sequenced on a PromethION flow cell. Illumina libraries were prepared using a modified Meyer & Kircher (2010) protocol with 250 bp fragmentation prior to HiSeq sequencing. Hi-C libraries were constructed following established chromatin crosslinking methods.

Whole-genome sequencing generated 112 GB of long-read data using the Oxford Nanopore PromethION platform (Novogene), supplemented by 66.37 GB of paired-end Illumina HiSeq data for base-level error correction. Chromosomal scaffolding was achieved through 104.11 GB of Hi-C sequencing data processed according to Belton et al. (2012). Transcriptomic annotation utilized 65.45 GB of RNA-seq data from multiple tissues.

A total of 389 hairtail specimens from Vietnam (n=151), China (n=223), and Korea (n=65) were preserved in 95% ethanol at 4°C (Fig. 2 B). We discovered a distinct fish species in the markets of Dandong (China) and Busan (South Korea) that displays pronounced morphological differences compared to locally harvested *T. japonicus*. We provisionally classify it as *Trichiurus* sp., distinguished by its notably larger yellowish eyes, yellowish-opaque dorsal fins, and an overall larger body size. In contrast, *T. japonicus* typically features smaller white-colored eyes, transparent dorsal fins, a smaller body size, and an oily epidermal texture. Although sellers asserted that these specimens were locally harvested, some local residents identified them as imported fish. They asserted that these specimens were locally harvested, but morphological and genetic analyses strongly suggested an Indian Ocean origin (See discussion). To investiagate further, we collected ten specimens of this fish (*Trichiurus* sp.) from each regional market for analysis. Morphological identification preceded molecular analysis using COX1 gene amplification primers: Forward: CTACAACCCACCGCTTACTC; Reverse: GTTGTAATAAAGTTAATGGCGC. PCR products from successfully amplified specimens (n=81) underwent Sanger sequencing and phylogenetic reconstruction via maximum likelihood analysis in MEGA-X with 100 bootstrap replicates (Kumar et al., 1994).

The study was initiated through COX1 gene amplification and sequencing of collected ribbon fish specimens. Subsequent phylogenetic tree construction and NCBI database comparisons revealed distinct taxonomic groupings among samples. Specimens collected from Vietnamese coastal waters were identified as *Trichiurus lepturus*. Samples obtained from Zhoushan, Xiamen, Qingdao, and Hainan regions were classified as *Trichiurus japonicus*. The *Trichiurus nanhaiensis* and *Trichiurus brevis* specimens were exclusively collected from Vietnam and Hainan. Notably, a taxonomically unresolved group from Korea and Northeast China showed highest BLAST similarity to *T. lepturus* in NCBI database, yet exhibited divergent phylogenetic positioning from *T. lepturus* in our tree topology.

The extracted DNA was used for library construction. DNA integrity was verified through NanoDrop 3300 fluorospectrometry and agarose gel electrophoresis. Ultrasonic fragmentation (Xinzhi Bio-Ultra DNA Fragmentator; Scientz) achieved consistent 250-1000 bp fragments using cyclic sonication parameters: 20s ON/20s OFF pulses at 300W over 22 cycles. Dual-indexed Illumina libraries incorporated inline barcodes during adapter ligation to prevent cross-contamination (Wang et al., 2022). Final library size selection (250-1000 bp) employed QIAquick Gel Extraction prior to HiSeq2500 sequencing.

To ensure population representation, five individuals from each potentially distinct geographic population were sequenced, ultimately establishing a resequencing library comprising 81 samples. An average sequencing depth of 8× coverage was achieved per specimen, with one optimal sample per species selected for deep sequencing at 40× coverage. Following stringent quality control procedures to exclude sequencing failures and contaminated samples, 57 high-quality specimens remained for subsequent genomic analyses. Post-QC exclusion of degraded or contaminated samples yielded high-quality resequencing data from 56 individuals.

### Genome assembly

Genome size estimation was performed using GenomeScope 2.0 (Vurture et al., 2017), a computational tool for analyzing k-mer frequency distributions. The 21-mer frequency profile was generated from Nanopore paired-end sequencing reads using Jellyfish v2.3.0 (Marcais and Kingsford, 2011), followed by genome size prediction through statistical modeling in GenomeScope2.0. De novo genome assembly was conducted using NextDenovo (Hu et al., 2024), a long-read assembler optimized for PacBio and Nanopore sequencing data, with default parameters and the estimated genome size as input. To improve assembly accuracy, a two-stage polishing strategy was implemented: 1) systematic error correction using NextPolish v1.4.1 (Hu et al., 2020) with Illumina short reads, followed by 2) haplotype reduction using Purge Haplotigs v1.1.2 (Roach et al., 2018) to eliminate redundant heterozygous sequences.

Hi-C data processing was performed using HiC-Pro v3.1.0 (Servant et al., 2015) with Bowtie2 v2.4.5 (Langmead and Salzberg, 2012) for sequence alignment and quality filtering. Chromosomal scaffolding was achieved through iterative processing with Juicer v1.6.2 (Durand et al., 2016) and 3D-DNA v180922 (Dudchenko et al., 2017), ultimately resolving 24 chromosome-scale scaffolds. Assembly visualization and manual curation were conducted using Juicebox v1.11.08 (Robinson et al., 2018; accessed June 2024). Genome completeness was assessed through BUSCO v5.4.3 (Simão et al., 2015) against the actinopterygii_odb10 database.

Repetitive elements were systematically identified using an integrative approach combining de novo prediction with RepeatModeler v2.0.3 (Flynn et al., 2020) and homology-based detection using RepeatMasker v4.1.2-p1 (Tarailo-Graovac and Chen, 2009). For gene structure annotation, a hybrid evidence-based pipeline was implemented: 1) Transcriptomic evidence from ten tissues was assembled using Trinity v2.13.2 (Grabherr et al., 2011) and refined with CD-HIT v4.8.1 (Fu et al., 2012); 2) Cross-species protein homology evidence was obtained from NCBI RefSeq entries of *Danio rerio* and related Scombriformes species; 3) Ab initio prediction was performed using MAKER3 v3.01.03 (Cantarel et al., 2008) with integrated training of Augustus v3.4.0 (Stanke et al., 2008) and SNAP v2013-11-29 (Korf, 2004) models.

Predicted genes underwent comprehensive functional annotation through sequential analysis: Homology searches using BLASTP v2.12.0+ (Camacho et al., 2009) against NCBI NR, KEGG, Pfam, and SwissProt databases; Domain architecture analysis via InterProScan v5.56-89.0 (Jones et al., 2014); Consensus annotation integration using evidence-based prioritization. Genome visualization was performed using pyCircos v1.0.1 (accessed June 2024), while synteny analysis with *Scomber japonicus* and *Thunnus albacares* was conducted using JCVI v1.1.14 (Tang et al., 2015) with default parameters.

### Resequencing and population genomic analysis

Raw sequencing data underwent demultiplexing using a dual-indexing strategy combining 8-bp i7 (P7) and i5 (P5) adapters with 6-bp sample-specific inline barcodes. Adapter contamination and low-quality sequences were removed through sequential processing with TrimGalore v0.6.7 (Krueger, 2020) and Cutadapt v1.2.1, implementing quality filtering (Phred score <20). A custom Perl pipeline (Yuan et al., 2019) was employed for read refinement and format standardization.

Single nucleotide polymorphism (SNP) discovery was performed following GATK v4.2.0.0 best practices (Poplin et al., 2018), including duplicate marking. Raw SNPs were filtered using GATK’s VariantFiltration with the following criteria: QUAL < 30.0, QD < 2.0, FS > 60.0, SOR > 4.0. Resultant variant call format (VCF) files were processed using PLINK v1.9 (Purcell et al., 2007) for data format conversion and quality control (genotype missing rate <0.2). Population structure analysis was conducted with ADMIXTURE v1.3.0 (Alexander et al., 2009) using cross-validation (K=1-7) to determine optimal ancestral clusters.

Pairwise sequentially Markovian coalescent (PSMC, Li et al., 2011) analysis was performed on whole-genome resequencing data from six representative species, each represented by a single diploid individual sequenced at 40× coverage. For species lacking existing genomic references, de novo draft genome assemblies were constructed using SOAPdenovo2 (Luo et al., 2012) with resequencing data achieving 40× sequencing depth per individual. Raw sequencing reads were processed through a standardized pipeline: adapter trimming using Cutadapt v3.4 with quality filtering (Q≥20), BWA-MEM v0.7.17 alignment to respective reference genomes, and 3) consensus sequence generation following established protocols. The PSMC analysis was implemented using the psmc v0.6.5 toolkit with default parameters (-p “4+25*2+4+6”). Generation time (g) was calculated as g = a + [s/(1-s)] = 7 years, where a = 3 years (adult maturation age from FishBase) and s = 0.5 (discrete generation overlap coefficient). Visualization of coalescent models employed psmc_plot.pl v1.0. To assess model robustness, 100 bootstrap replicates per species were performed through random segment sampling followed by trajectory smoothing using a cubic spline interpolation.

The Assexon methodology (Yuan et al., 2019), originally designed for exon capture sequencing data assembly and phylogeny reconstruction, adapting it for whole-genome resequencing analysis given current cost reductions in sequencing technologies. The analytical pipeline requires three primary inputs: (1) paired-end resequencing data, (2) conserved exon sequences (both amino acid and nucleotide levels) and reference genome data from a closely related species. For this study, we utilized 4,434 annotated genes from *Oreochromis niloticus* and its genome as the reference dataset. Data were processed by: PCR duplicates removal, UBLAST-aligned sorting (ubxandp.pl), SGA-based de novo assembly, Exonerate-filtered exons, reciprocal BLAST validation (reblast.pl), yielding reference-aligned consensus sequences. MAFFT-aligned loci were converted to PHYLIP (concat_loci.pl) for RAxML (A. Stamataki, 2014) based phylogeny. phylogenetic reconstruction. This dual approach enables simultaneous evaluation of phylogenetic relationships while accounting for incomplete lineage sorting through multi-locus coalescent methods.

Following completion of the aforementioned procedures, rigorous validation was required to detect potential contamination or sequencing errors in the initial mitochondrial and COI sequences, thereby ensuring taxonomic reliability of species assignments. For this purpose, resequencing data were subjected to mitochondrial genome assembly using the GetOrganelle pipeline (Jin et al., 2020). Samples successfully generating circularized and complete mitochondrial genomes were subsequently annotated through the Mitos platform (M. Bernt et al., 2013). The resultant COX1 gene sequences were then extracted and aligned for phylogenetic tree reconstruction. This multi-stage analytical approach enabled systematic verification of species identities through evaluation of evolutionary relationships within the established phylogenetic framework.

The species delimitation analysis for *Trichiurus* spp. was conducted using FastSimCoal v2.8 (Excoffier, L et al., 2021). Building upon previously established phylogenetic relationships, multiple alternative models representing different speciation sequences were constructed, with the (*lepturus*-*japonicus*, *japonicus*-*nanhaiensis*, *nanhaiensis*-sp.) model emerging as statistically optimal through comparative scoring analysis. Using this best-supported model, three key evolutionary parameters were simulated: inter-species gene flow patterns, divergence timing (initial generation range: 200-300,000 generations), and effective population sizes across lineages. The analysis incorporated coalescent simulations to account for ancestral population structure and stochastic variance in genealogical outcomes.

To investigate genomic signatures of positive selection in *Trichiurus japonicus*, a selective sweep analysis was conducted using the SweeD (Pavlidis et al., 2013) software. A total of 20 individuals were sampled to capture genetic diversity. For selective sweep detection, SweeD v3.3.2 was employed under default parameters, utilizing a composite likelihood ratio (CLR) framework to identify loci deviating from neutral expectations. The analysis was performed on a chromosome-wide basis, with likelihood scores calculated in non-overlapping 10-kb windows. Genomic regions under selection were defined as those within the genomic regions with CLR scores exceeding the 99th percentile (p < 0.01). Genes located within these selected loci were functionally annotated and comparatively analyzed.

## Author contribution

VG, TW, and CL conceived the research plan. VG collected the samples, TW and VG did experiments and collected data. TW analyzed the whole data and make the figures. LL, JH and MZ contributed to data analysis. LL, JH and MZ participated in experimental work. TW and CL wrote the draft. All authors edited and approved the final version of the manuscript.

## Acknowledgment

We extend special thanks to Prof. Hanlin Wu for encouraging the research team to address taxonomic complexities in *Trichiurus*. We also acknowledge Dr. Shaoxiong Ding, Dr. Hui Wang, Mr. Jiajie Chen, and Ms. Ni Xin for their assistance with specimen collection.

## Funding

This work was supported by the Science and Technology Commission of Shanghai Municipality (19410740500; 19050501900) to CL.

## Data availability

The scaffold-level genome assemblies of the *Trichiurus japonicus* (JBITNA000000000; BioSample SAMN44500129) have been deposited in NCBI (https://www.ncbi.nlm.nih.gov/) genome database. The Chromium-linked Illumina reads, RNA-Seq reads (Illumina HiSeq), and Nanopore reads used for the genome assembly have been deposited to NCBI genome database (SRA) under the BioProject accession number PRJNA1179659.

All data have been deposited in the Dryad repository under the URL: http://datadryad.org/share/GpQ6KcJfWqSEUvww62AUhzWXb-arxI_ZrA9TvkG45Us. All other relevant data are available from the corresponding authors upon request.

## Conflict of interest

the authors declare that there is no conflict of interest.

## Notes

### Competing Interest Statement

The authors have declared no competing interest.

### Summary of Updates

In our initial study, we collected Trichiurus specimens from commercial markets in Dandong, China and Busan, South Korea, where local vendors consistently identified them as locallocal hairtail species. Based on this locality information and preliminary morphological assessments, we previously posited the existence of a monophyletic T. sp population within the Yellow Sea ecosystem. However, comprehensive molecular analyses conducted in this revised study, including mitochondrial DNA cytochrome b sequencing and comparative phylogenetic reconstruction with Indo-Pacific reference sequences, demonstrate that the specimens cluster within the Indian Ocean Trichiurus lineage. This phylogeographic discrepancy likely reflects undocumented transoceanic trade networks supplying processed fisheries products to Northeast Asian markets. Consequently, all previous conclusions regarding Yellow Sea population structure, species-specific ecological adaptations, and regional fisheries management implications have been retracted. The revised manuscript incorporates corrected distribution maps, updated discussion of Indo-Pacific Trichiurus dispersal mechanisms, and caveats regarding specimen provenance verification in commercial catch studies. This revision underscores the critical importance of integrating molecular validation with catch documentation in marine biogeographic research.

## Reference

[1] Abdul-Muneer, P. M. (2014). Application of microsatellite markers in conservation genetics and fisheries management: recent advances in population structure analysis and conservation strategies. Genetics research international, 2014(1), 691759.

[2] Alexander, D.H., J. Novembre, and K. Lange, Fast model-based estimation of ancestry in unrelated individuals. Genome Res, 2009.

[3] A. Stamatakis: “RAxML Version 8: A tool for Phylogenetic Analysis and Post-Analysis of Large Phylogenies”. In Bioinformatics, 2014

[4] Belton JM, McCord RP, Gibcus JH, Naumova N, Zhan Y, Dekker J. Hi-C: a comprehensive technique to capture the conformation of genomes. Methods. 2012 Nov;58(3):268–76. doi: 10.1016/j.ymeth.2012.05.001.

[5] Bouckaert, R., Heled, J., Kühnert, D., Vaughan, T., Wu, C-H., Xie, D., Suchard, MA., Rambaut, A., & Drummond, A. J. (2014). BEAST 2: A Software Platform for Bayesian Evolutionary Analysis. PLoS Computational Biology, 10(4), e1003537. doi:10.1371/journal.pcbi.1003537

[6] Camacho, C., Coulouris, G., Avagyan, V., Ma, N., Papadopoulos, J., Bealer, K., & Madden, T. L. (2009). BLAST+: architecture and applications. BMC bioinformatics, 10, 1–9.

[7] Cantarel, BL; Korf, I; Robb, SMC;, et al. MAKER: An easy-to-use annotation pipeline designed for emerging model [J].GENOME RESEARCH, 2008. DOI: 10.1101/gr.6743907.

[8] Chakraborty, A., Aranishi, F., & Iwatsuki, Y. (2006). Genetic differentiation of Trichiurus japonicus and T. lepturus (Perciformes: Trichiuridae) based on mitochondrial DNA analysis. ZOOLOGICAL STUDIES-TAIPEI-, 45(3), 419.

[9] Chu, Y.T. (1931). Index Piscium Sinensium. Biological Bulletin of St. John’s University, Shanghai,1, 1–290

[10] Dudchenko O, Batra S S, Omer A D, et al. De novo assembly of the Aedes aegypti genome using Hi-C yields chromosome-length scaffolds[J]. Science, 2017, 356(6333): 92–95.

[11] Durand, N., Shamim, M., Machol, I., Rao, S. P., Huntley, M., Lander, E., & Aiden, E. L. (2016). Juicer Provides a One-Click System for Analyzing Loop-Resolution Hi-C Experiments. [J]. Cell Systems, 3, 95–98.

[12] Excoffier, L., Marchi, N., Marques, D. A., Gouy, A., Sousa, V. C. (2021) fastsimcoal2: demographic inference under complex evolutionary scenarios. Bioinformatics 37: 4882 4885.

[13] FAO. (2021). Fishery and Aquaculture Statistics (2020 statistical yearbook). Food and Agriculture Organization of the United Nations.

[14] Flynn, JM; Hubley, R; Goubert, C; Rosen, J; Clark, AG; Feschotte, C; Smit, AF. RepeatModeler2 for automated genomic discovery of transposable element families. PNAS. DOI: 10.1073/pnas.1921046117.

[15] Forskål P. (1775). Descriptiones Animalium, Avium, Amphibiorum, Piscium, Insectorum, Vermium; quae in Itinere Orientali Observavit Petrus Forskål. Post Mortem Auctoris editit Carsten Niebuhr. Adjuncta est materia Medica Kahirina. Mölleri, Hafniae, 19 + xxxiv + 164 pp.

[16] Goode, G. B., & Bean, T. H. (1882). Benthodesmus, a new genus of deep-sea fishes, allied to Lepidopus. Proceedings of the United States National Museum.

[17] Golani, D., & Fricke, R. (2018). Checklist of the Red Sea fishes with delineation of the Gulf of Suez, Gulf of Aqaba, endemism and Lessepsian migrants. Zootaxa, 4509(1), 1–215.

[18] Grabherr, MG; Haas, BJ; Yassour, M; Levin, JZ;, et al. Full-length transcriptome assembly from RNA-Seq data without a reference genome [J]. NATURE BIOTECHNOLOGY 2011. DOI: 10.1038/nbt.1883.

[19] Guo, Y. H., Halasan, L. C., Wang, H. Y., & Lin, H. C. (2022). High migratory propensity constitutes a single stock of an exploited cutlassfish species in the Northwest Pacific: A microsatellite approach. Plos one, 17(3), e0265548.

[20] Hermann, von, & Wissmann. (2002). The pleistocene glaciation in china*. Bulletin of the Geological Society of China.

[21] Hong-Liang, H., Feng-Hua, T., Xue-Zhong, C., Zhang, H., Ling-Zhi, L., Xue-Feng, S., … & De-Hu, W. (2016). Nets selectivity of capsule size diamond mesh of Trichiurus haumela in East China Sea during Summer. Journal of Agriculture Resources and Environment, 33(5), 433.

[22] Hu, J; Fan, JP; Sun, ZY; Liu, SL. NextPolish: a fast and efficient genome polishing tool for long-read assembly. [J]. BIOINFORMATICS 2020. DOI: 10.1093/bioinformatics/btz891.

[23] Hu, J; Wang, Z; Sun, ZY; Hu, BX; Ayoola, AO; Liang, F; et al. NextDenovo: an efficient error correction and accurate assembly tool for noisy long reads. [J].GENOME BIOLOGY 2024. DOI: 10.1186/s13059-024-03252-4.

[24] Javier Gago, F. (1997). Character evolution and phylogeny of the cutlassfishes: an ontogenetic perspective (Scombroidei: Trichiuridae). Bulletin of marine science, 60(1), 161–191.

[25] Jiao Lishi. Molecular Phylogenetics and Feeding Habits of East China Sea Hairtail (*Trichiurus* spp.) [D]. Zhejiang Ocean University, 2024. DOI:10.27747/d.cnki.gzjhy.2024.000083.

[26] Jin, J. J., Yu, W. B., Yang, J. B., Song, Y., DePamphilis, C. W., Yi, T. S., & Li, D. Z. (2020). GetOrganelle: a fast and versatile toolkit for accurate de novo assembly of organelle genomes. Genome biology, 21, 1–31.

[27] Jin, X., Shan, X., Li, X., Wang, J., Cui, Y., & Zuo, T. (2013). Long-term changes in the fishery ecosystem structure of Laizhou Bay, China. Science China Earth Sciences, 56, 366–374.

[28] Jones, P., Binns, D., Chang, H. Y., Fraser, M., Li, W., McAnulla, C., … Hunter, S. (2014). InterProScan 5: genome-scale protein function classification. Bioinformatics, 30, 1236–1240.

[29] Kim, Y. H., Yoo, J. T., Lee, E. H., Oh, T. Y., & Lee, D. W. (2011). Age and growth of largehead hairtail Trichiurus lepturus in the East China Sea. Korean Journal of Fisheries and Aquatic Sciences, 44(6), 695–700.

[30] Krueger F. Trim galore. Wrapper Tool Cutadapt FastQC Consistently Apply Qual Adapt Trimming FastQ Files, 2015, 516. https://github.com/FelixKrueger/TrimGalore (15 March 2024, date last accessed).

[31] Krueger F, Andrews SR. Bismark: a flexible aligner and methylation caller for Bisulfite-Seq applications. Bioinformatics 2011;27:1571–2.

[32] Kumar, S., Tamura, K., & Nei, M. (1994). MEGA: molecular evolutionary genetics analysis software for microcomputers. Bioinformatics, 10(2), 189–191.

[33] Langmead, B., Trapnell, C., Pop, M., & Salzberg, S. L. (2009). Ultrafast and memory-efficient alignment of short DNA sequences to the human genome. Genome Biology, 10, R25.

[34] Li A. et al. DNA Barcoding Identification and Distribution of Sand Hairtail (L.sasala) in the Yellow Sea. Journal of Fishery Sciences of China, 2022, 29(05): 696–703.

[35] Li, C. (2006). Discussion on the original records and type specimens of *Trichiurus lepturus* Linnaeus 1758. In Chinese Society of Zoology & Chinese Society of Oceanology and Limnology, Ichthyology Branch (Eds.), Abstracts of the 7th Congress of the Ichthyology Branch and Symposium Celebrating the 110th Anniversary of Professor Chu Y.T.’s Birth (p. 1). Institute of Oceanology, Chinese Academy of Sciences. (Chinese version)

[36] Li, S. (1947). The Lushan Ice Age. Shanghai: Commercial Press.

[37] Li C.S. (1992). “Hairtail fishes from Chinese coastal waters (Trichiuridae).” Mar Sci Acad Sin 4: 212–218.

[38] Li H., Durbin R. (2011). Inference of human population history from individual whole-genome sequences. Nature. 475, 493–496. doi: 10.1038/nature10231

[39] Lin, H. C., Tsai, C. J., & Wang, H. Y. (2021). Variation in global distribution, population structures, and demographic history for four *Trichiurus* cutlassfishes. PeerJ, 9, e12639.

[40] Linnaeus, C. (1758). Systema Naturae per regna tria naturae, secundum classes, ordines, genera, species, cum characteribus, differentiis, synonymis, locis. Editio decima, reformata [10th revised edition], vol. 1: 824 pp. Laurentius Salvius: Holmiae.

[41] Liu, Y., Cheng, J. H., & Jia, S. G. (2021). Characteristics of submarine water temperature distribution of Trichiurus haumela in the East China sea and Southern Yellow sea with the improvement of the analysis methods.

[42] Luo, R., Liu, B., Xie, Y., Li, Z., Huang, W., Yuan, J., … & Wang, J. (2012). SOAPdenovo2: an empirically improved memory-efficient short-read de novo assembler. Gigascience, 1(1), 2047–217X.

[43] Martins, A. S., & Haimovici, M. (1997). Distribution, abundance and biological interactions of the cutlassfish *Trichiurus lepturus* in the southern Brazil subtropical convergence ecosystem. Fisheries research, 30(3), 217–227.

[44] Meyer, M. & Kircher, M. 2010. Illumina sequencing library preparation for highly multiplexed target capture and sequencing. Cold Spring Harbor Protocols, 2010, pdb. prot5448.

[45] Mirarab, S., Reaz, R., Bayzid, M. S., Zimmermann, T., Swenson, M. S., & Warnow, T. (2014). ASTRAL: genome-scale coalescent-based species tree estimation. Bioinformatics, 30(17), i541–i548.

[46] Miya, M., Friedman, M., Satoh, T. P., Takeshima, H., Sado, T., Iwasaki, W., Yamanoue, Y., Nakatani, M., Mabuchi, K., Inoue, J. G., Poulsen, J. Y., Fukunaga, T., Sato, Y., & Nishida, M. (2013). Evolutionary origin of the Scombridae (tunas and mackerels): members of a paleogene adaptive radiation with 14 other pelagic fish families. PloS one, 8(9), e73535.

[47] Mohammed, R. S., Ramjohn, C., Lucas, F., & Rostant, W. G. (2010). Additional observations on the distribution of some freshwater fish of Trinidad and the record of an exotic. Living World, Journal of the Trinidad and Tobago Field Naturalists’ Club, 43–53.

[48] Mthethwa S, Bester-van der Merwe AE, Roodt-Wilding R. Addressing the complex phylogenetic relationship of the Gempylidae fishes using mitogenome data. Ecol Evol. 2023 Jun 21;13(6):e10217. doi: 10.1002/ece3.10217.

[49] Mukundan, L. P., Sukumaran, S., Sebastian, W., & Gopalakrishnan, A. (2020). Characterization of the whole mitogenome of largehead hairtail Trichiurus lepturus (Trichiuridae): Insights into special characteristics. Biochemical Genetics, 58, 430–451.

[50] M. Bernt, A. Donath, F. Jühling, F. Externbrink, C. Florentz, G. Fritzsch, J. Pütz, M. Middendorf, P. F. Stadler MITOS: Improved de novo Metazoan Mitochondrial Genome Annotation Molecular Phylogenetics and Evolution 2013, 69(2):313–319

[51] Nakabo T. 2002. Family Trichiuridae. In T Nakabo, ed. Fishes of Japan with pictorial keys to the species. Tokyo: Tokai Univ. Press, p. 1142. (in Japanese)

[52] Okamoto, N., Koike, K., Shibata, J. Y., & Tomiyama, T. (2018). Species composition of hairtails (Trichiuridae) in Myanmar. Regional Studies in Marine Science, 17, 73–77.

[53] Panhwar, S. K., Dong, Z. Y., Tianxiang, G., Ping, W., Zhiqiang, H., Zhongming, W., & Yang, W. (2018). Decadal Population Traits and Fishery of Largehead Hairtail, Trichiurus lepturus (Linnaeus, 1758) in the East China Sea. Pakistan Journal of Zoology, 50(1).

[54] Pavlidis, P; Zivkovic, D; Stamatakis, A; Alachiotis, N. SweeD: Likelihood-Based Detection of Selective Sweeps in Thousands of Genomes. [J] MOLECULAR BIOLOGY AND EVOLUTION DOI:10.1093/molbev/mst112.

[55] Poplin, R., Ruano-Rubio, V., DePristo, M. A., Fennell, T. J., Carneiro, M. O., Van der Auwera, G. A., … & Banks, E. (2017). Scaling accurate genetic variant discovery to tens of thousands of samples. BioRxiv, 201178.

[56] Purcell, S., Neale, B., Todd-Brown, K., Thomas, L., Ferreira, M. A., Bender, D., Maller, J., Sklar, P., de Bakker, P. I., Daly, M. J., & Sham, P. C. (2007). PLINK: A tool set for whole-genome association and population-based linkage analyses. The American Journal of Human Genetics, 81(3), 559–575

[57] Roach, M. J., Schmidt, S. A., & Borneman, A. R. (2018). Purge Haplotigs: Allelic contig reassignment for third-gen diploid genome assemblies. BMC Bioinformatics, 19(1), 1–10

[58] Robinson, J. T., Turner, D., Durand, N. C., Thorvaldsdóttir, H., Mesirov, J. P., & Aiden, E. L. (2018). Juicebox. js provides a cloud-based visualization system for Hi-C data. Cell systems, 6(2), 256–258.

[59] Servant, N., Varoquaux, N., Lajoie, B. R., Viara, E., Chen, C. J., Vert, J. P., Heard, E., Dekker, J., & Barillot, E.(2015). HiC-Pro: an optimized and flexible pipeline for Hi-C data processing. Genome Biology, 16, 259.

[60] Simão, F. A., Waterhouse, R. M., Ioannidis, P., Kriventseva, E. V., & Zdobnov, E. M. (2015). BUSCO: Assessing genome assembly and annotation completeness with single-copy orthologs. Bioinformatics, 31(19), 3210–3212.

[61] Stanke, M; Keller, O; Gunduz, I; Hayes, A; Waack, S; Morgenstern, B.AUGUSTUS:: ab initio prediction of alternative transcripts [J]. NUCLEIC ACIDS RESEARCH, 2008. DOI: 10.1093/nar/gkl200.

[62] Tang, H., Krishnakumar, V., Zeng, X., Xu, Z., Taranto, A., Lomas, J. S., … & Zhang, X. (2024). JCVI: A versatile toolkit for comparative genomics analysis. iMeta, e211.

[63] Tarailo-Graovac, M., & Chen, N. (2009). Using RepeatMasker to identify repetitive elements in genomic sequences. Current Protocols in Bioinformatics, Chapter 4, Unit 4.10.

[64] Temminck CJ, Schlegel H (1843) Pisces. Part 1. In: von Siebold, PF (ed) Fauna Japonica. Müller, Amsterdam, pp 1–20

[65] Tzeng, C. H., & Chiu, T. S. (2012). DNA barcode-based identification of commercially caught cutlassfishes (Family: Trichiuridae) with a phylogenetic assessment. Fisheries Research, 127, 176–181.

[66] Tzeng, C. H., Tsan-Yu, C. H. I. U., Chih-Shin, C. H. E. N., Hui-Yu, W. A. N. G., & Tai-Sheng, C. H. I. U. (2016). The Current Population Structure of the Demersal Hairtail (Trichiurus japonicus) in the Western North Pacific was Shaped by the Taiwan Strait, as Revealed by Mitochondrial DNA. Taiwania, 61(4).

[67] Vurture, GW; Sedlazeck, FJ; Nattestad, M; Underwood, CJ; Fang, H; Gurtowski, J; Schatz, MC. GenomeScope: fast reference-free genome profiling from short reads [J]. BIOINFORMATICS 2017. DOI:10.1093/bioinformatics/btx153.

[68] Wang Keling, Zhang Peijun, Liu Lanying, et al. Study on Species Differentiation of Hairtail (Trichiuridae) in China’s Coastal Waters. Acta Oceanologica Sinica (Chinese Edition), 1993, (02): 77–83+145-146.

[69] Wang, K.-L.; Liu, L.-Y.; You, F.; Xu, C. (1992). Studies on the genetic variation and systematics of the hairtalls [hairtail] fishes from the South China Sea. Marine Science (Beijing). 2: 69–72

[70] Wang, K., You, F., Xu, C., et al. (1995). Commentary on “Hairtail fishes from Chinese coastal waters (Trichiuridae)”. Oceanologia et Limnologia Sinica, 26(2), 215–222

[71] Wang, Ying., Yuan, H., Huang, J. & Li, C. 2022b. Inline index helped in cleaning up data contamination generated during library preparation and the subsequent steps. Molecular Biology Reports, 1–8.

[72] Wang, H. Y., Dong, C. A., & Lin, H. C. (2017). DNA barcoding of fisheries catch to reveal composition and distribution of cutlassfishes along the Taiwan coast. Fisheries Research, 187, 103–109.

[73] Wen Hui. Genetic Diversity of Japanese Hairtail (*Trichiurus japonicus*) along China’s Coast and Comparative Analysis of Mitochondrial Genomes among Three Hairtail Species in the South China Sea [D]. Guangdong Ocean University, 2023. DOI:10.27788/d.cnki.ggdhy.2023.000219.

[74] Tokimura M, Yamada U, Trie T. Comments on taxonomy and distributions of the species of Thrichiurus in the East China and Yellow Seas and adjacent waters[J]. Seikaiku Suisan Kenkyusho News, 1995, 80: 12–14.

[75] Ye, Y., & Rosenberg, A. A. (1991). A study of the dynamics and management of the hairtail fishery, Trichiurus haumela, in the East China Sea. Aquatic Living Resources, 4(2), 65–75.

[76] Yi M-R, Hsu K-C, Gu S, He X-B, Luo Z-S, Lin H-D, Yan Y-R (2022) Complete mitogenomes of four *Trichiurus* species: A taxonomic review of the *T. lepturus* species complex. ZooKeys 1084: 1–26.

[77] Yuan, H., Atta, C., Tornabene, L. & Li, C. 2019. Assexon: assembling exon using gene capture data. Evolutionary Bioinformatics, 15, 1176934319874792.

[78] Z. Yang and B. Rannala. Bayesian estimation of species divergence times under a molecular clock using multiple fossil calibrations with soft bounds. Mol Biol Evol, 23(1):212–26, 2006.

